# Cancer Cells Viscoelasticity Measurement by Quantitative Phase and Flow Stress Induction

**DOI:** 10.1101/2021.08.05.455201

**Authors:** Tomas Vicar, Jaromir Gumulec, Jiri Chmelik, Jiri Navratil, Radim Kolar, Larisa Chmelikova, Vratislav Cmiel, Ivo Provaznik, Michal Masarik

## Abstract

Cell viscoelastic properties are affected by the cell cycle, differentiation, pathological processes such as malignant transformation. Therefore, evaluation of the mechanical properties of the cells proved to be an approach to obtaining information on the functional state of the cells. Most of the currently used methods for cell mechanophenotypisation are limited by low robustness or the need for highly expert operation. In this paper, the system and method for viscoelasticity measurement using shear stress induction by fluid flow is described and tested. Quantitative Phase Imaging (QPI) is used for image acquisition because this technique enables to quantify optical path length delays introduced by the sample, thus providing a label-free objective measure of morphology and dynamics. Viscosity and elasticity determination were refined using a new approach based on the linear system model and parametric deconvolution. The proposed method allows high-throughput measurements during live cell experiments and even through a time-lapse, where we demonstrated the possibility of simultaneous extraction of shear modulus, viscosity, cell morphology, and QPI-derived cell parameters like circularity or cell mass. Additionally, the proposed method provides a simple approach to measure cell refractive index with the same setup, which is required for reliable cell height measurement with QPI, an essential parameter for viscoelasticity calculation. Reliability of the proposed viscoelasticity measurement system was tested in several experiments including cell types of different Young/shear modulus and treatment with cytochalasin D or docetaxel, and an agreement with atomic force microscopy was observed. The applicability of the proposed approach was also confirmed by a time-lapse experiment with cytochalasin D washout, where an increase of stiffness corresponded to actin repolymerisation in time.

**SIGNIFICANCE:** We present an approach for viscoelasticity measurement using QPI and shear stress induction by fluid flow. Our system builds and extends a recently published approach by parametric deconvolution, which allows us to eliminate the influence of the fluidic system and reliably measure both the shear modulus and viscosity of the cells in high throughput. Additionally, the proposed method enables to simultaneously determine cell refractive index map, cell dry mass map, and morphology, thereby enabling a multimodal cellular characterisation in a single measurement.

## INTRODUCTION

The mechanical response of cells is affected by the cell cycle, differentiation, pathological processes such as malignant transformation, cardiovascular diseases, or aging (1–3). Evaluation of the mechanical properties of the cells thereby proved to be an approach to obtaining information on the functional state of the cells potentially relevant for the clinical setting (4).

There are plenty of methods of cell mechanical characteristics measurements based on various principles, including Atomic Force Microscopy (AFM), microfluidics, micropipette aspiration, magnetic or optical tweezers, particle tracking rheology, and acoustic methods (5, 6). However, the measured values are not comparable between those methods with variations in the mechanical properties of cells by orders of magnitude (7, 8).

A widely established method for cell mechanophenotypisation is AFM (9, 10). This method is based on direct contact of a cantilever tip, from the degree of its deflection Young modulus is calculated. As the cantilever mechanically scans the cells with a single probe, it non-physiologically stresses the cells, which can unnaturally affect the results. Furthermore, the method is also low-throughput and limited by the necessity of highly expert operators and the need for the selection of appropriate probes (11). Microfluidic-based approaches can on the other hand be implemented in a high-throughput setup. These methods employ various strategies including micro-constriction of cells in small channels, flow in cross-slot microchannels, or application of shear forces. With an appropriate model, these approaches can obtain more information on the mechanical properties (viscosity and elasticity). The connection of flow shear stress induction and Quantitative Phase Imaging (QPI), introduced by Eldrige et al. (12, 13), brings a new possibility to reliably evaluate the viscoelastic properties of the cells. Compared to common light microscopy techniques, QPI enables rapid determination of the cell geometry, including the estimation of cell height, a parameter needed for shear modulus estimation.

The systems for shear stress experiments typically consist of flow chambers, which allow the generation of stable shear stress gradients using a suitable pump and tubing. The characteristics of each component determine the final properties of the entire system, which might have a significant effect on the subsequent analysis of the cell’s properties. This can be especially important in cases where the responses to the fast temporal changes of the shear stress are examined. To minimize the influence of the fluidic system, a deconvolution-based approach for the estimation of the cell mechanical properties was employed.

To estimate the shear modulus, the cell height needs to be estimated. Although this can be calculated from the phase and cell refractive index, the latter is typically considered a constant for the whole cell population. Further refinement was made possible by estimation of refractive index using a method published by Rappaz et al. (14). In this approach, two perfusion media with different refractive indexes are used to estimate both cell refractive index and cell height.

In this paper, we describe a complex QPI-based approach for shear modulus estimation consisting of hardware and software parts including simultaneous refractive index determination and parametric deconvolution. The proposed method allows fast and reliable measurements during live-cell experiments, where QPI images can be used for simultaneous analysis of cell morphology.

## 1 MATERIALS AND METHODS

### 1.1 Hardware setup

The block scheme of our experimental setup with photos of the individual components is shown in Figure 1. A programmable dual syringe pump (Syringe pump Chemyx Fusion 4000) is utilized as a flow source. The chamber used in our setup is a commercially available, shear-stress optimized chamber (Ibidi *μ*-Slide VI 0.1). The flow meter (Sensirion SLF3S-1300F) is connected to our setup at the output of the flow chamber. All components are connected to appropriate adapters and tubing. To ensure the rigidity of the system, we have selected the PTFE material of the tubing. Our preliminary experiments showed that the main influence on the flow temporal profile has the syringe inserted in the syringe pump and the tubing material as shown in Supplementary Figure 8.

**Figure 1:**
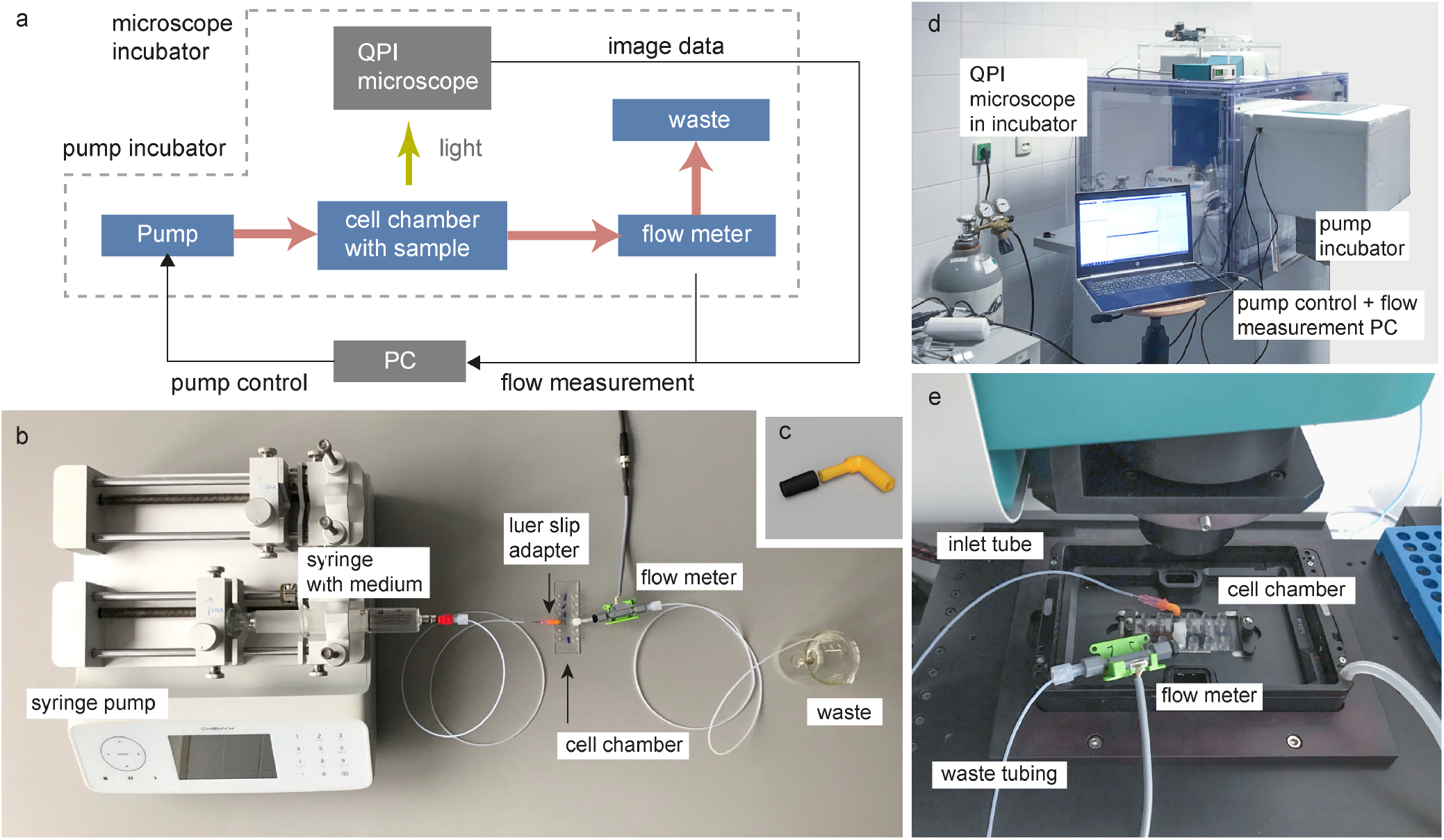
Shear flow system. (A) Block diagram of components; pink arrows indicate medium flow and the yellow arrow indicates light passes through the sample. (B) Photo of connected components. (C) Detail of 3D model of custom luer slip to μ-Slide VI 0.1 adapter. The black part is a rubber plastic preventing fluid leakage. (D) Photo of whole fluidic setup. (E) Detail of cell chamber inside the microscope.

For the acquisition of Quantitative Phase Images (QPI), a coherence-controlled holographic microscope (Telight, Q-Phase) was used. Objective Nikon Plan 10×/0.3 was used for hologram acquisition with a CCD camera (XIMEA MR4021MC). Holographic images were numerically reconstructed with the Fourier transform method (described in (15)) and phase unwrapping was used on the phase image. Moreover, background compensation with the polynomial fitting was applied, which set the background phase delay to zero. Near-maximum available frame rate (3 frames/s) was used to acquire cell movements with the highest temporal resolution.

Shear stress *τ*_0_ for the flow rate *Q* provided by syringe pump was calculated based on Ibidi μ-Slide IV 0.1 Application Note (16), which provides the same values as the calculation 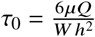 from (13), where *μ* is the dynamic viscosity of the medium, *W* is the width of the flow channel, and *h* is the height of the flow channel. At this scale, the drag force of the flowing liquid is negligible compared to the shear stress. The value of actual flow in the system was measured by a liquid flow sensor connected to the output of the flow chamber (see Figure 1).

### 1.2 Data processing pipeline

The whole data processing pipeline is shown on Figure 2a, where image sequence and measured flow are processed to estimate viscoelastic properties of cells, specifically, shear modulus *G* and viscosity *η*. First, the shear stress signal *τ(t)* is calculated from the measured flow. Second, individual cells or cell clusters are segmented and tracked in the acquired image stack and their center of mass movement over time (*CoM(t)*) is extracted. Moreover, thanks to the quantitative property of the QPI image, the cell height *L*_*cell*_ is estimated from the QPI image. Shear strain signal *γ(t)* can then be estimated from *CoM(t)* and *L*_*cell*_, as shown in Figure 2b. Finally, the viscoelastic properties (*G* and *η*) are estimated from the shear stress and shear strain signal using parametric deconvolution. A more detailed explanation of the individual data processing steps follows.

**Figure 2:**
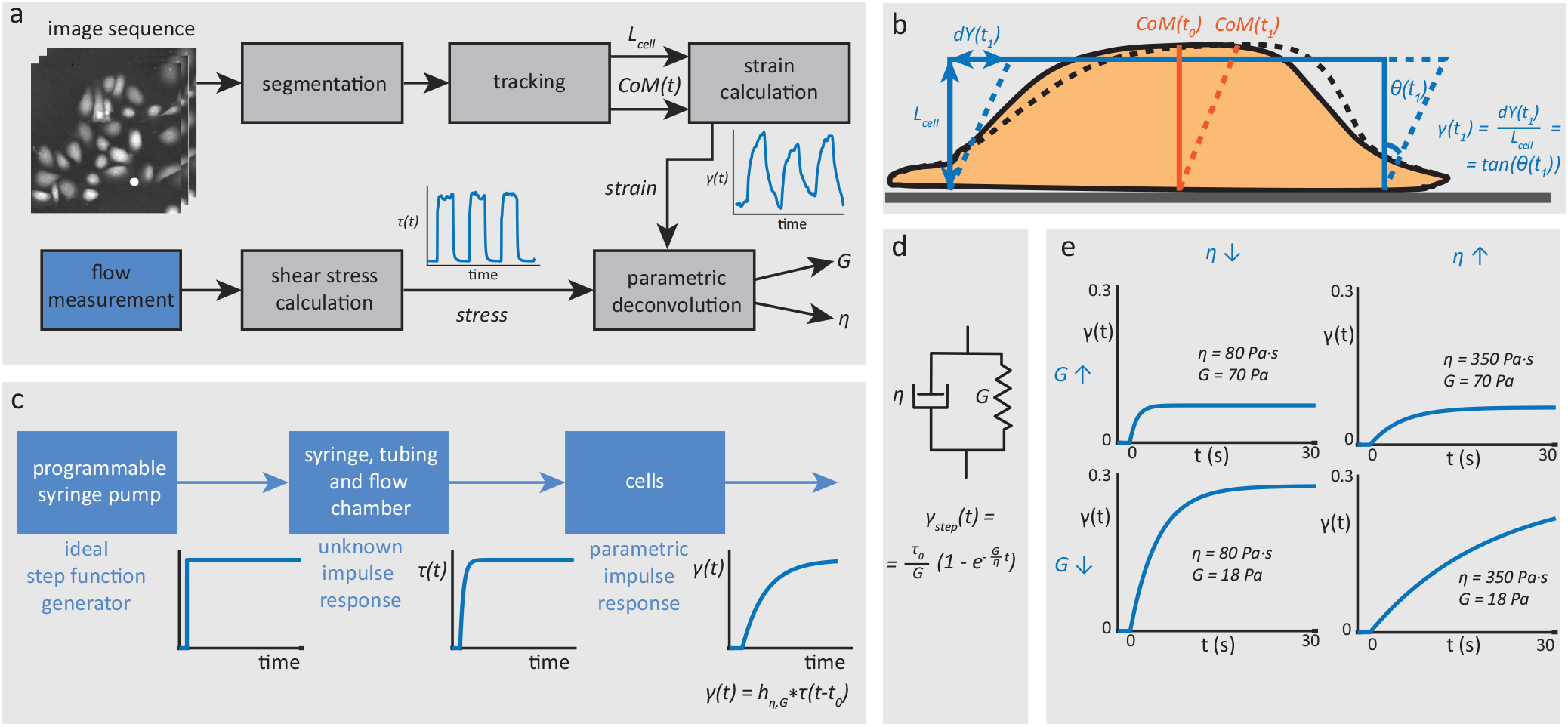
Data processing and viscoelasticity measurement explanation. (A) Block diagram of the data processing pipeline. (B) A model used in our calculation - the cell is approximated by a block with specific dimensions (solid lines); the block is skewed during the flow (dashed lines). (C) Strain signal generation model for parametric deconvolution based on a model of the connected linear systems. (D) Schematic representation of Kelvin-Voight (KV) model, which can be represented by the parallel combination of a purely viscous damper and purely elastic spring. (E) Example of shear strain response to ideal step function for low and high viscosity/shear modulus.

### 1.3 Image processing

To measure the viscoelastic properties of the cells, images are processed to get the height of individual cells and temporal changes of centroid position (*CoM(t)*). Cell height and *CoM(t)* are used for the calculation of the shear strain signal, which is consequently used for the calculation of viscoelastic parameters as described in Section 1.4. The individual steps needed to obtain shear strain for a particular cell or cell cluster are described below. From here on, the analysis is described for cells, however, the same applies to the cell clusters. Despite the analysis of cell clusters provides a single number for whole cluster (average of viscoelastic parameters of cells in cluster), it is more accurate in scenarios with touching cells which lead to segmentation errors. This results in fluctuation of cell borders in time, thereby causing noisy shear strain signals.

#### Segmentation

Stack of reconstructed phase video frames was filtered by a 3D median filter (of size 3 × 3 × 7 for *x*, *y*, *t*, respectively) to suppress noise in each frame and to filter the areas, where the phase values are distorted due to fluid flow. The segmented cells were obtained by thresholding of the filtered image using a small positive threshold value of 0.35 rad, generally corresponding to the lowest phase values on the cell periphery. Obtained binary frames were post-processed by area filtering, which removed objects smaller than the selected threshold of 100 px (39.27 *μm*^2^). Cells touching the border of the actual field of view were removed in each frame to achieve the correct *CoM(t)* estimation.

#### Tracking

Due to the 3D median filtering used in the segmentation step and the relatively high acquisition frame rate, the segmented cells create a compact 3D object in the video stack, i.e., the cell area in each frame is overlapped with the same cell area in the following frame. When each separate cell in the first video frame got a unique label, and the 3D connected component analysis was done through all frames starting from the first one, the same cell obtained the same label through the whole video sequence. In the post-processing step, the cell was removed from further analysis if: 1) the cell was not presented in the first frame (probably false detection such as a bubble or part of a dead or detached cell), and/or 2) the cell was not detected in at least 60% of all frames (e.g., not well-adhered cell). Finally, the position of the center of mass of each cell was computed in each video frame as a weighted average of the values of the mass density image belonging to the segmented cell.

#### Centre of Mass Movement

The Centre of Mass *CoM(t)* signal for particular cell is extracted from the reconstructed phase image *ϕ*(*x*, *y*). CoM movement of each cell was computed as the difference between the actual *CoM*(*t*) and initial *CoM*(*t*_0_), where the movement was considered in the direction of the applied flow only (in the y-axis only from the image point of view), which produces signal *dY(t)* used in the shear strain calculation as shown on Figure 2b. The movement in the direction orthogonal to the applied flow (x-axis) could be caused only by unwanted cell migration in our experiments.

#### Cell Height Measurement

QPI microscope provides the phase image *ϕ(x, y)* that represents the phase delay produced by the cell, which depends on the difference of refractive indexes (RIs) between the cells and the surrounding medium. Assuming a homogeneous cell with constant refractive index, the cell height map can be estimated as (17):

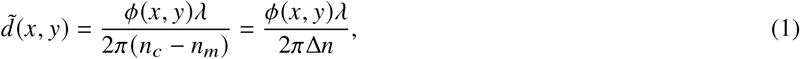

where *λ* = 650*nm* is light wavelength and *n*_*c*_ and *n*_*m*_ is RI of cells and surrounding medium, respectively. The RIs estimation is described in Section 1.5. Height of the individual cells was calculated as the median of cell pixels over the whole time stack (*L*_*cell*_). The *L*_*cell*_ value represents the cell height required for shear strain calculation.

#### Shear Strain

Finally, the shear strain temporal change can be calculated from CoM movement signal *dY(t)* and cell height *L*_*cell*_ as:

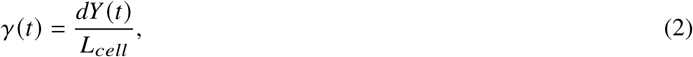

which is schematically depicted on Figure 2b. In this model, the cell is approximated with a rectangular block of of homogeneous material of constant viscosity and shear modulus, which are estimated using this method.

### 1.4 Viscoelastic model estimation

A Kelvin-Voight (KV) model is a linear model often used for the representation of viscoelastic properties of a single cell (18). The parallel connection of spring (with elastic modulus G, N/m) and a damper (with viscosity *η*, Ns/m) enables to quantify the cell elasticity and viscosity (see Figure 2d). When this model is applied in a setup involving shear stress, the quantity G is referred to as a shear modulus and it expresses the stiffness of the particular cell. Furthermore, for isotropic elasticity, the linear relation between the shear modulus and Young’s modulus exists (13).

The response of KV system to the positive step change of the external shear stress can be described by an exponential function as:

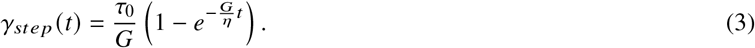

The quantity *τ*_0_ represents the magnitude of the shear stress step function, which is set during the experiments. Examples of shear strain response to step function shear stress with various *G* and *η* are shown in Figure 2e. These parameters can be obtained by an exponential fitting of the cell response signal represented by the equation 2. As mentioned in Section 1.1, the non-rigid properties of the entire setup influence the whole experiment, particularly the shear stress signal *τ(t)* and shear strain *γ(t)* signals. This influence can be modeled as a serial connection of linear systems as shown in Figure 2c. The input of this connection is a signal driving the movement of the syringe. This is typically a step function or a square signal. The generated flow (i.e., the shear stress) is distorted by the flow system, which can be described by an unknown impulse response.

Consequently, the distorted shear stress waveform *τ*(*t* – *t*_0_) deforms the adhered cells and change their *CoM*(*t*). Under the assumption of linear systems, the final model of the shear strain *γ*_*model*_(*t*) is a convolution:

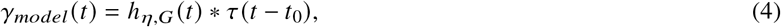

where *h*_*η*,*G*_*(t)* is a parametric impulse response with unknown parameters *η* (cell viscosity) and *G* (cell shear modulus), *τ(t – t_0_)* is a shear stress applied on the cell (generally shifted by unknown shift *t*_0_). The parametric impulse response *h*_*η*,*G*_*(t)* can be easily computed as a differentiation of the step response (eq. 3):

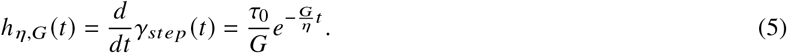

This convolution model enables to formulate a simple cost function: ∥*γ(t) – γ*_*model*_*(t)*∥_1_, or more specifically ∥*γ(t) – τ*(*t* – *t*_0_) * *h*_*η*,*G*_*(t)*∥_1_, and estimate the parameters of KV model using suitable optimization method. Parametric deconvolution is described in more details in (19).

### 1.5 Cell height calculation and refractive index measurement

The cell height calculation according to the equation 1 is not straightforward as the exact RI values (*n*_*c*_ and *n*_*m*_) are not known. The refractive index of the medium *n*_*m*_ can be easily measured by a refractometer. However, the cell RI *n*_*c*_ is unknown.

Fortunately, the presented setup of shear stress induction enables to relatively easily estimate *n*_*c*_. Specifically, two syringes with medium of different (but known) RIs can be connected to the cell chamber, and they can be used for fast exchange of the surrounding medium (see Figure 3a). Thus, we can measure image *ϕ*_1_(*x*, *y*) with the first medium with RI *n*_*m*1_ (Figure 3b – upper left) and image *ϕ*_2_(*x*, *y*) with the second medium with RI *n*_*m*2_ (Figure 3b – upper right). Therefore, if we assume that the cell height *d*(*x*, *y*) and RI of cell *n*_*cell*_ are equal for both measurements, then:

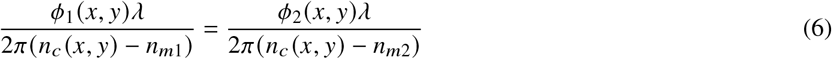

**Figure 3:**
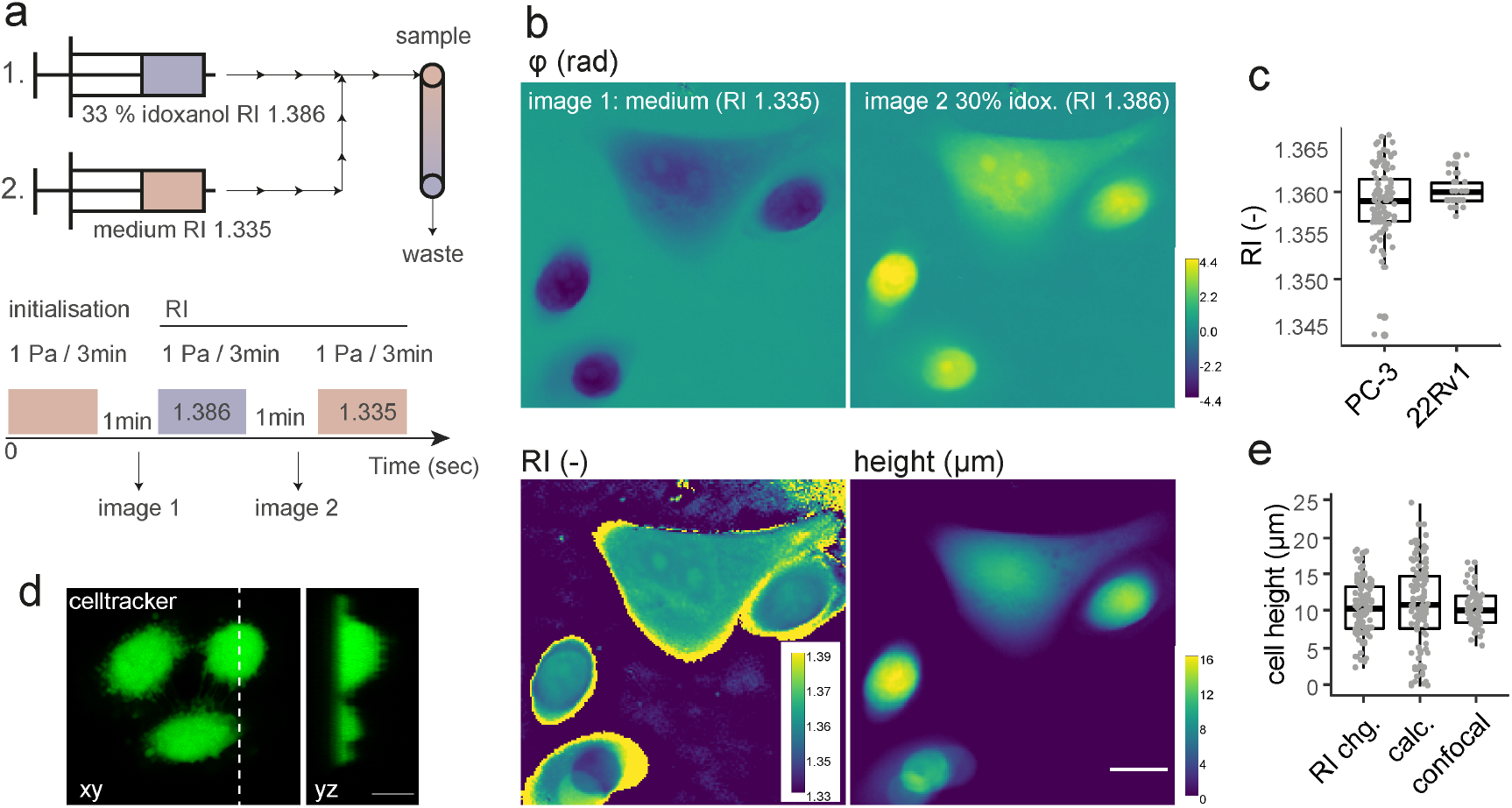
Refractive index measurements. (A) RI measurement setup diagram of medium exchange. (B) Images acquired with different media solutions (idoxanol solution and mixture of idoxanol and RPMI medium); cell RI image was calculated with the equation 7, and cell height image was calculated with equation 8. (C) Measured RIs for PC-3 and 22Rv1 cells – values are medians of manually selected cell interior of RI image (D) An example of confocal microscopy image used for cell height verification. (E) Comparison of cell height calculated by equation 8, equation 1 and measured by confocal microscopy. Scalebar correspond to 10 *μm*.

By rearrangement of the terms in this equation, the RI of cells can be calculated as:

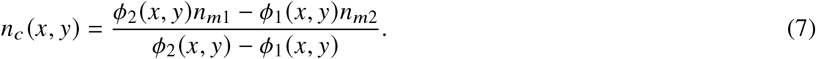

This equation can be used to compute the RI for each pixel and to estimate the RI of a specific cell line as an average of RI values inside these cells (Figure 3b – lower left). Moreover, this approach enables to compute the spatial distribution of cell height (i.e., for each pixel position) using a combination of equation 1 and 7:

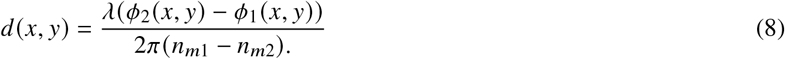

Application of this double-syringe approach provides a possibility to measure the cell height *d(x, y)* for every pixel in the field of view (FOV) in the shear induction experiments (Figure 3b – lower right). However, this requires a medium exchange measurement for each measured FOV, which is time-consuming and might not be necessary for morphologically homogeneous cell populations. Thus, to simplify the measurements, RI (i.e., *n*_*c*_) for each cell line was determined only once using the equation 7 and this value was used in equation 1 to determine the cell height. Cell height of individual cells for shear modulus calculation in this paper was calculated as the median of cell height image 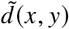 of the cell over the whole image sequence based on the equation 1.

### 1.6 Flow sequence

First, a 60 s initialisation pulse is applied with flow 129.8 *μ/min*. This flow caused shear stress of 1 Pa, which minimally influenced the cells, as tested in the optimisation. The purpose of initialisation is to fill the outlet tubing with medium, flush out non-adherent cells and debris, and remove microbubbles. After this initialization, a sequence of 3 × 30 s pulses of 5 Pa shear stress with 30 s pauses between them was used, as shown on Figure 4a. Responses to the whole sequence serve as input to parametric deconvolution viscoelasticity estimation, where the purpose of repeated pulses is to make the estimation more precise and robust.

**Figure 4:**
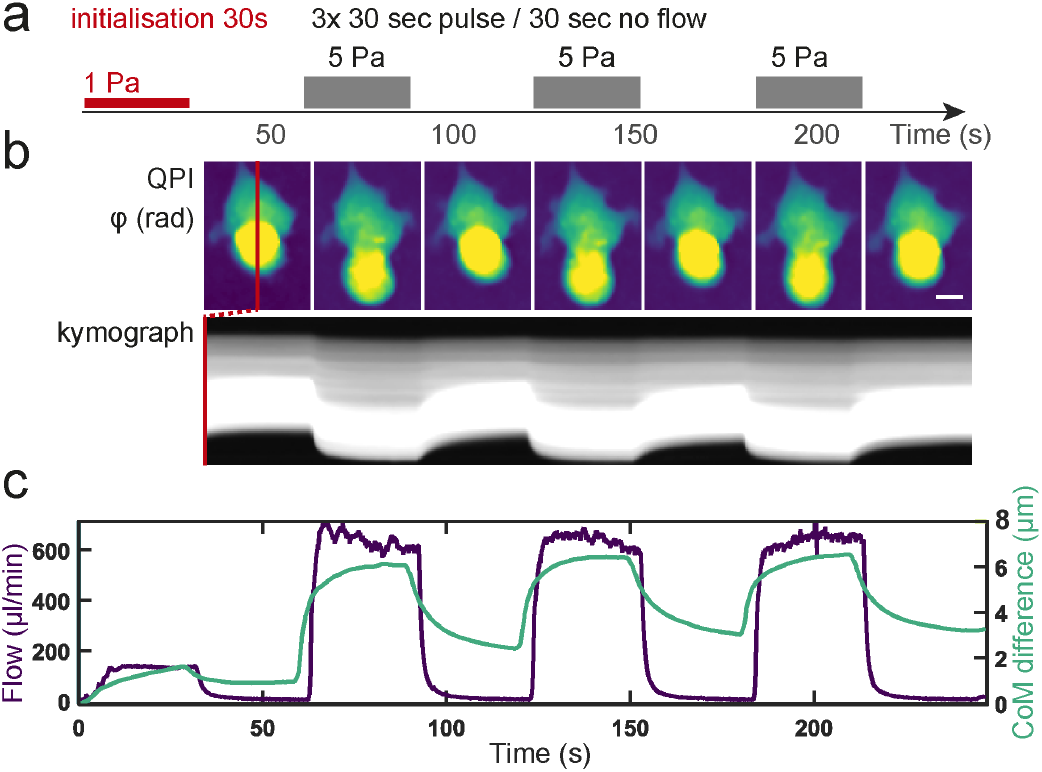
Cell response to standardized shear stress sequence. A. Schematic of standardized experiment shear stress sequence. B. kymograph of representative 22Rv1 cell cluster. C. plot showing measured flow and cell centre of mass (CoM) difference over flow sequence. Scalebar corresponds to 10 *μm*.

## 2 RESULTS

### 2.1 Refractive index

As described in Section 1.5, the measurement of cell RIs is required to reliably estimate the cell height. Calculation of RI image is possible by the acquisition of images of cells with two different media of two different RIs. The first medium *m*1 was the RPMI medium (RI 1.3353) used in shear stress induction experiments, and the second medium *m*2 consists of 33.4 % idoxanol (RI 1.3864), which was demonstrated to be suitable for live cell experiments (20). Example of images with *m*1 and *m*2, and the calculated RI and cell height image using equation 7 and 8 are shown in Figure 3b.

Measured RI for PC-3 and 22Rv1 cells is shown in Figure 3c. Values are medians of manually labeled cell interiors, in order to omit large noisy values on cell edges. The estimated height of the cells was confirmed by measurement using a confocal microscope (example on Figure 3d). A box plot of this comparison is shown in Figure 3e – similar medians of cell heights were measured using confocal microscopy, using the calculated cell height image *d(x, y)* (equation 8) and cell height calculated from the phase image using the measured RI (equation 1), which is used for our viscoelasticity analysis.

### 2.2 Cell types

Cell lines previously used in our lab (10) characterised by different Young modulus, were used to test the system. PC-3 and 22Rv1 cells were used as a model of low- and highly-deformable cells, respectively. Cell culture conditions are described in Supplementary material 4.1. The standardized shear stress sequence was applied – Figure 5a, where shear strain value optimalisation is available in Supplementary material 4.4.

**Figure 5:**
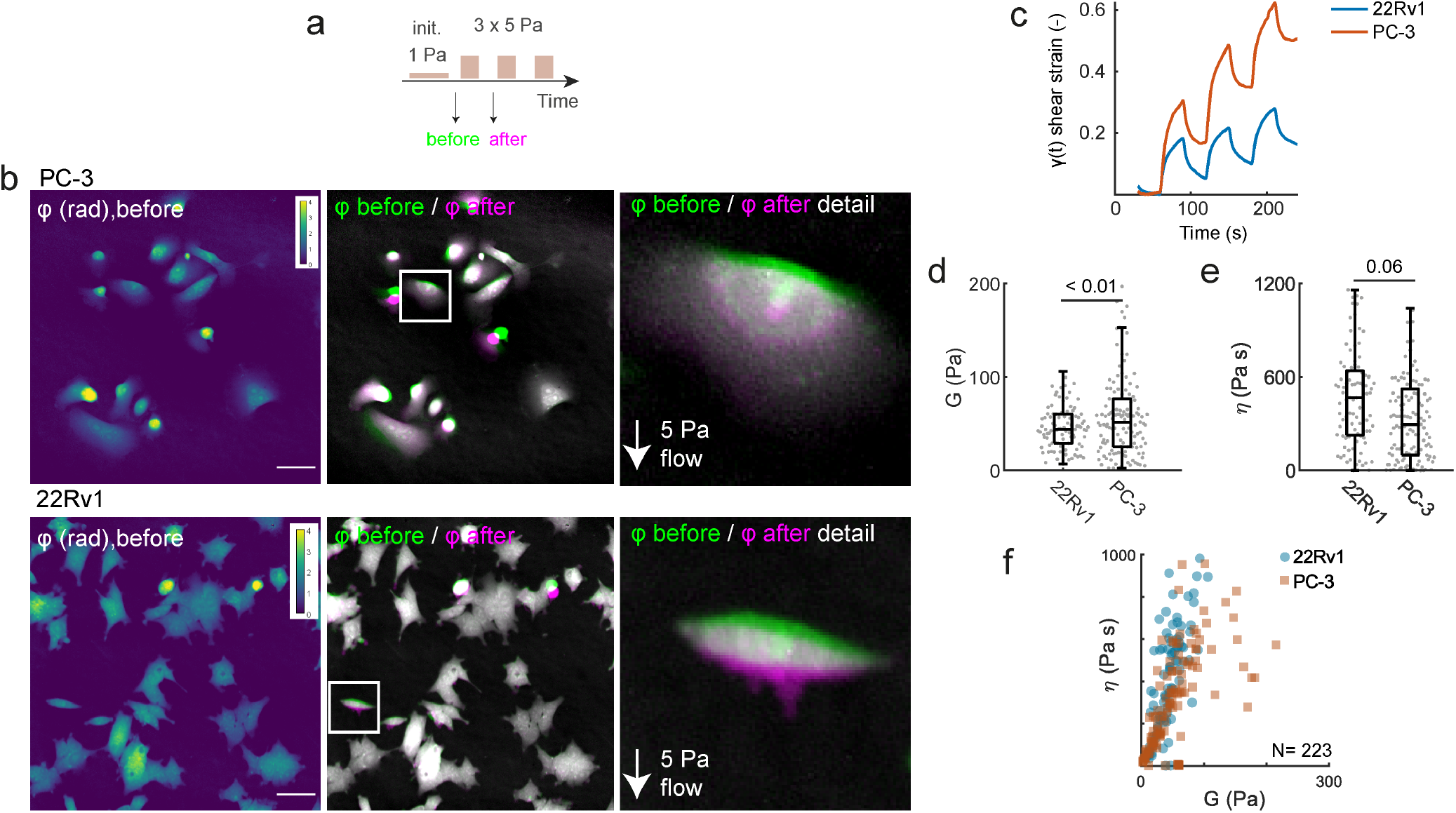
Results of viscoelasticity measurement of different cell types (22Rv1 and PC-3). (A) Schematics of applied shear stress sequence. (B) Example of images before applied shear stress (left), a merge of the image before and with applied shear stress (middle), and zoom of this merge (right). (C) The plot of average strain signal of PC-3 and 22Rv1 cells. (D) Results of shear modulus. (E) Results of viscosity. (F) Scatter plot of shear modulus and viscosity. Scalebar indicates 50 *μ*m.

Example images with applied shear stress are shown on Figure 5b. We have determined how the cells differ in a stress-strain response Figure 5c and we have extracted viscoelastic parameters using parametric deconvolution. Resulting shear modulus shown an agreement of stiffness trend with previously published results, i.e. that PC-3 cells have higher stiffness than 22Rv1 cells (see Figure 5d). Apart from *G*, also viscosity *η* is presented on Figure 5e. However, compared to *G* the 22Rv1 cells have higher viscosity – 468.7 Pa·s for 22Rv1 vs. 296.5 Pa·s for PC-3. Figure 5f shows the dependence of viscosity and shear modulus for these two cell types, where values have linear relation, however, the relation has different slopes for these two cell types. This difference can be also seen from the average shear strain signal on Figure 5c: response of 22Rv1 is faster due to smaller elasticity, but it finally achieved a similar strain as PC-3 due to lower viscosity.

### 2.3 Treatments

Similarly, the responses of PC-3 cells to two types of treatments were tested to prove the validity of the proposed viscoelasticity measurements, which were measured again with our standardized experiment shear stress sequence (Figure 5a) and analyzed using parametric deconvolution. Cells were affected by the cytoskeleton-targeting drugs Cytochalasin D (CytD) and docetaxel. The average shear strain signal response is shown on Figure 6b. Specifically, CytD is known to interfere with actin polymerisation (21) and was applied to decrease the cell stiffness and cytoplasmatic viscosity (22), and docetaxel was applied to increase cell stiffness based on (23). For details about the treatment dose and application, see Supplementary material 4.1.

**Figure 6:**
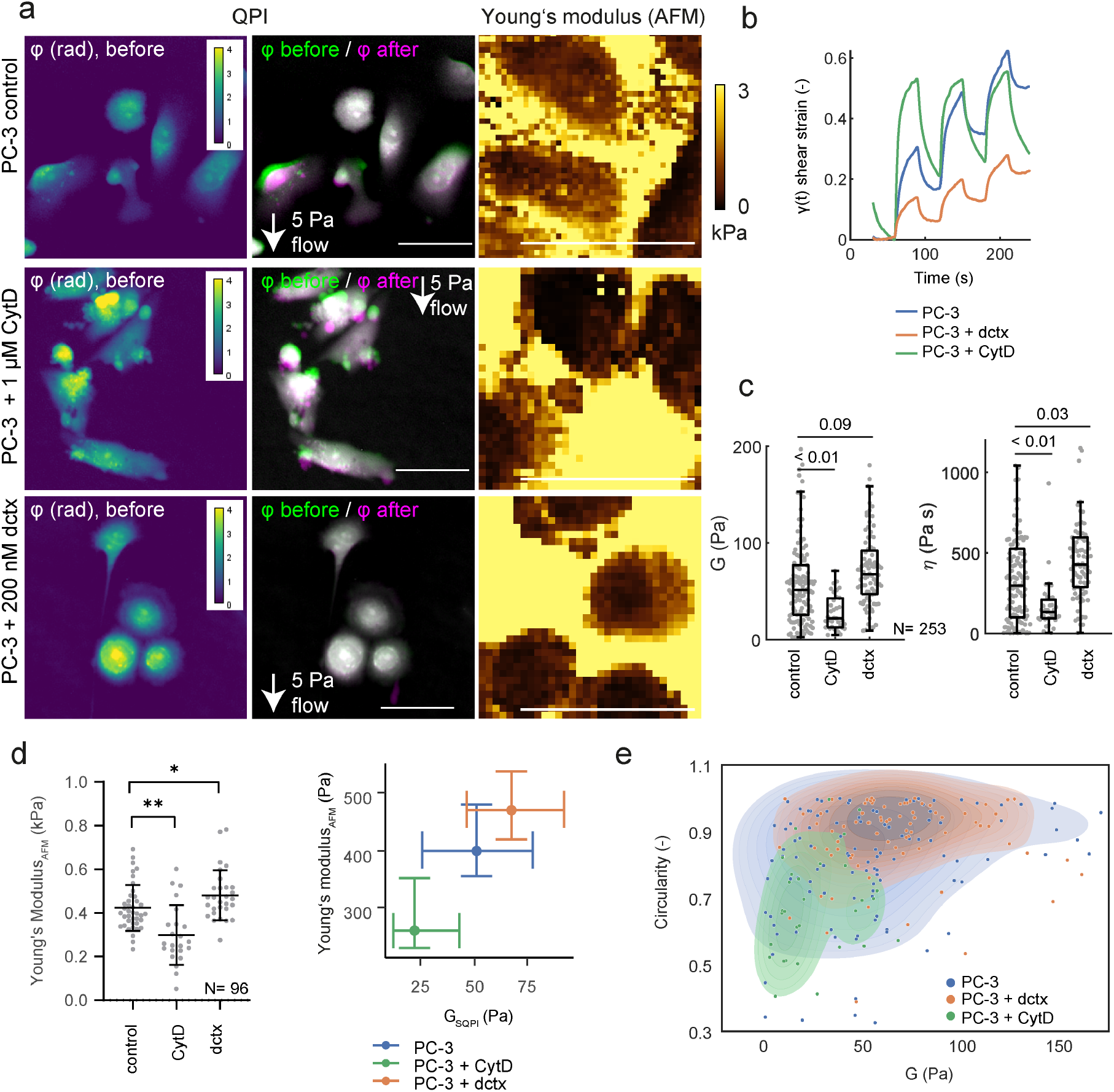
Results of viscoelasticity measurement of PC-3 cells treated with Cytochalasin D (CytD) and docetaxel (dctx) compared with AFM measurement. (A) An example of images before applied shear stress (left), the merge of the image before and with applied shear stress (middle), and AFM measured Young’s modulus example (right). (B) Plot of average strain signal for individual treatments. (C) Results of shear modulus and viscosity. (D) Atomic force microscopy measurement of Young modulus (left, total 92 cells), and a scatterplot of Young modulus determined by AFM a QPI. (E) Scatter plot of shear modulus and cell circularity, outlines correspond to kernel density estimate. Scalebar indicates 50 *μ*m.

Results of the measured shear modulus and viscosity are shown in Figure 6c. The measured stiffness was, as expected, higher for docetaxel treatment and lower for CytD treatment with 51.4 Pa, 22.0 Pa and 67.5 Pa for untreated, docetaxel treated and CytD-treated cells, respectively. Viscosity followed a similar trend with 296.2 Pa·s, 132.9 Pa·s and 427.7 Pa·s for untreated, docetaxel treated and CytD treated cells, respectively. The acquired data were furthermore verified by AFM (Figure 6d), where the shear modulus values were roughly 9fold lower compared to Young modulus and cell elastic parameters determined by both methods demonstrated to be linearly dependent. For AFM methodology, see Supplementary material 4.2.

As a consequence of CytD-mediated inhibition of actin polymerisation the cells change also their morphology, where membrane blebbing was evident Figure 6a. Docetaxel-mediated tubulin stabilisation on the other hand results in a tubulin rearrangement which can also be reflected in cell morphology (10). Cell morphological parameters can be extracted from the acquired images so dependence between morphology and viscoelastic properties of the cells can be studied using the proposed approach. Circularity, as shown in Figure 6e, demonstrated to be different between those treatments. For the evaluation of circularity, only individual cells (not cell clusters) were manually selected, where the circularity was calculated as in (24). Data indicate that a combination of viscoelastic and morphological parameters can be beneficial to differentiate cell populations, when compared to differentiation based just on viscosity or elasticity. Similarly, parameters calculated from phase-cell dry mass, dry mass density (25), or refractive index can further supplement such analyses for more complex cell characterisation.

### 2.4 Actin cytoskeleton recovery

To demonstrate the possibility to measure in a time-lapse setting, we measured viscoelasticity during 20 minutes of actin repolymerisation after washout of CytD. We have observed that the morphology of CytD treated cells recovered quickly after application of flow during shear stress induction experiments (see example image on Figure 7b). As measured in (26), full recovery of actin fibers of most cell compartments is expected within 15-30 min after CytD washout. To measure cell viscoelasticity during actin recovery, we have designed a longer shear stress induction sequence consisting of 4 standardized sub-sequences of 3 pulses as shown in Figure 7a, where average shear strain response signal for individual sub-sequences at different times is shown on Figure 7c, where reduction of shear strain deviation is observable as cells recover from CytD treatment. Each sub-sequence is used in parametric deconvolution to measure viscoelasticity. As shown in Figure 7d–f, elasticity, viscosity, and circularity have recovered to almost nontreated values within 16 minutes after CytD washout. For the evaluation of circularity, only individual cells (not cell clusters) were manually selected, where the circularity was calculated as in (24).

**Figure 7:**
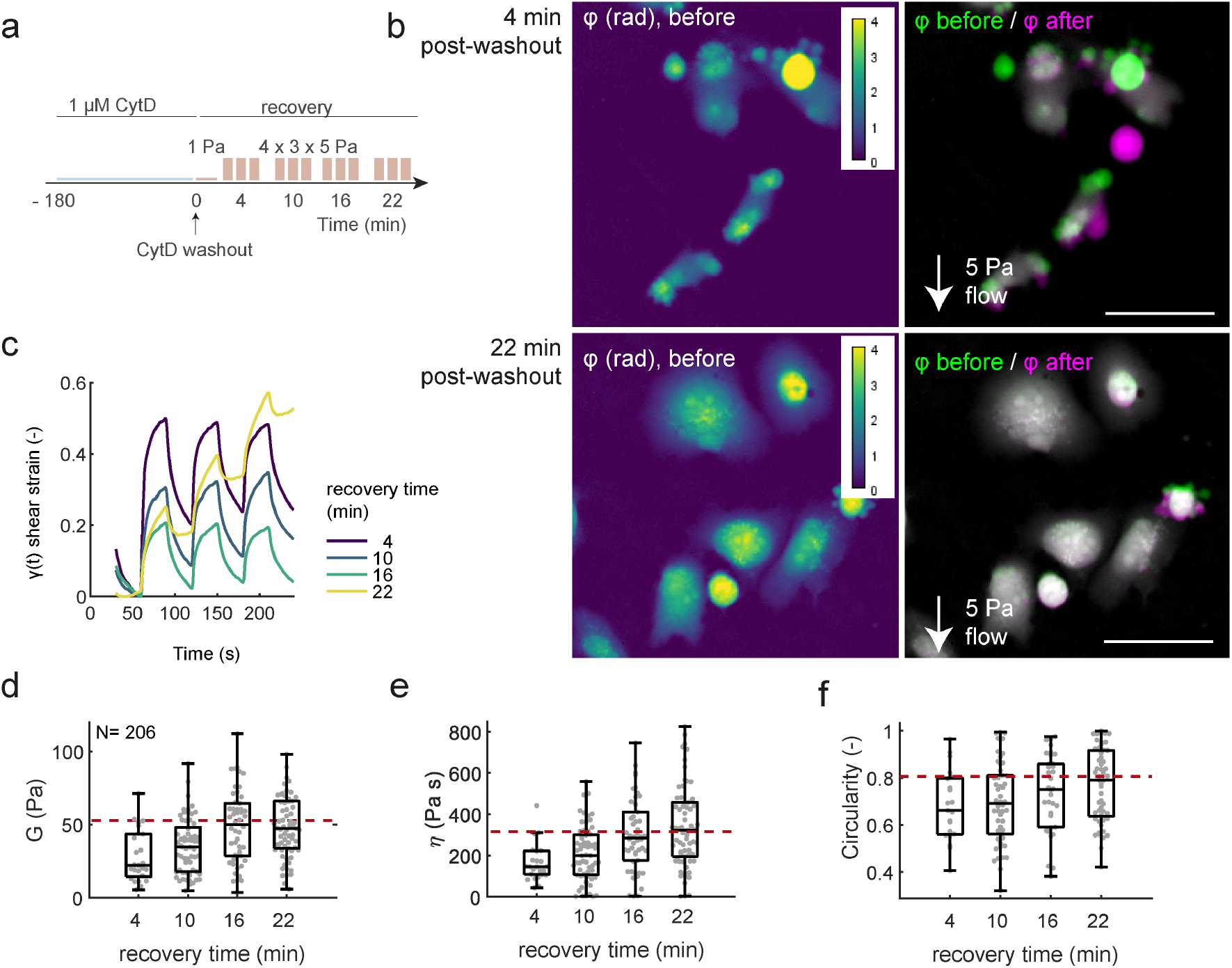
Results of viscoelasticity measurement of PC-3 cells after Cytochalasin D (CytD) washout. (A) Schematics of applied shear stress sequence. (B) An example of images acquired in 4^*th*^ and 22^*nd*^ minute during CytD washout (left) and example of its reaction to the shear stress – merge of the image before and with applied shear stress (right). (C) The plot of average strain signals during CytD washout recovery. (D-F) Results of shear modulus, viscosity, and cell circularity during CytD washout recovery. Red dashed line indicates median modulus, elasticity and circularity values for untreated cells. Scalebar indicates 50 *μ*m

## 3 DISCUSSION

Evaluation of the mechanical properties of the cells proved to be an approach to obtaining information on the functional state of the cells. Most of the currently used methods for cell mechanophenotypisation are limited by low robustness or the need for highly expert operation. In this paper, the system and methodology for viscoelasticity measurement using quantitative phase imaging (QPI) and shear stress induction by fluid flow are described and tested. We have tested the whole system in different types of experiments to show its benefits and properties. The proposed system and method are based on the original approach of Eldrige et al. (13); however, our approach includes also a reliable method to measure the refractive index directly during the measurement. RI of measured cells or cell clusters is then used to determine the cell height from the phase image. This is crucial as this value strongly influences the precision of model parameters estimation. There are two possibilities how to determine the cell height. The more time-consuming approach applies two perfusion media before each experiment to measure two phase images and to determine the cell height map. The final map provides a spatial distribution of the cell height over the whole field-of-view. During this procedure, the determined RI image map can be correlatively used with other phase-derived imaging data like cell dry mass. It has been demonstrated that RI is relevant for cellular processes like cell cycle and an important parameter for pathologies, including cancer and hematological diseases (27). The second possibility is to change the perfusion media only one time for a specific set of experiments to determine an average cell RI.

Additionally, the proposed estimation also provides the cell viscosity. Its value, however, is largely influenced by the system components – especially the tubing and syringe used in the pump. We, therefore, demonstrated that the issue is diminished by the parametric deconvolution and the viscosity values are then independent of the flow system characteristics (see characteristics of various setups in Supplementary Figure 8). Elimination of the effect of the system provides also practical benefits for the measurement setup. The plastic syringes are significantly easier to manipulate with, and unlike glass syringes, their pistons seal properly. Moreover, the applied pulses are not that steep, which makes it more gentle to the cells, thus, fewer cells are detached by flow during the experiment. Parametric deconvolution also provides the possibility to apply different flow waveforms (not only rectangular pulses) to estimate cell viscoelastic properties. Furthermore, the elimination of the flow system properties via deconvolution provides a way to compare model parameters between laboratories.

In agreement with our previous study (10), we confirmed that PC-3 cells measured by the proposed method have larger stiffness than 22Rv1 cells; however, PC-3 cells have rather lower viscosity, which shows that viscosity measurement can provide additional information that can be biologically interesting (28). The reliability of the proposed viscoelasticity measurement was also confirmed by force mapping using atomic force microscopy. Shear modulus showed a similar trend to Young modulus in cells exposed to docetaxel and cytochalasin D, respectively. Following cytochalasin treatment, a similar trend in elasticity was observed in Wakatsuki (21) study and specifically, an agreement regarding shear modulus was described in Eldridge et al. (13) study. In accordance with Spedden et al. (29) study, an overall increase in global cell stiffness was observed following treatment with paclitaxel, another taxane-based drug, using AFM-based technique.

Compared to AFM, this approach is easy to implement on a QPI microscope. In addition, the proposed approach is able to measure a large number of cells quickly as it measures the whole field of view of cells simultaneously (10-50 cells using our QPI microscope field of view size). Moreover, this hands-on approach is applicable during live-cell experiments, while QPI images can be used for simultaneous analysis of cell morphology, which can provide interesting information like cell viability or the distinction between apoptosis and necrosis (25). It also allows for multiple measurements of viscoelasticity during the experiment, which e.g., allows monitoring of influence of specific treatment. We have shown this in the CytD washout experiment, where we have performed multiple viscoelasticity measurements in time, while simultaneously observing cell morphology.

Apart from the Young modulus, viscosity is also important for understanding cell physiology. It is evidenced that it is relevant for cell status evaluation and cell-type classification (30) and it is a suitable indicator for *in vitro* evaluation of drug effects on cells (31). Increase in viscosity after cell division was described by Adeniba et al. (32) study due to cortical stiffening around mitosis in MCF-7 cells using a non-contact interferometric measurement technique. Reports indicate that viscosity is relevant also for cancer cell classification. Shimolina and colleagues described the involvement of membrane viscosity in the acquisition of cancer drug resistance (33) and interestingly, a study dating to 1941 reports tumor cells to be characteristic by a higher viscosity compared as determined by high-speed centrifugation (34). However, compared to cell elasticity, cell viscosity still remains not a well understood area. Regarding the effect of cytoskeleton-targeting drugs on viscosity, our results are in agreement with Wang et al. (30) study who demonstrated a viscosity decrease in H1299 cells following CytD treatment by micropipette-based measurement. Conversely to our results, Yun et al. (31), determined that the factor of cell viscosity, a parameter characterizing the relative viscous/elastic behavior of materials under external stresses, decreased in HeLa cells exposed to docetaxel. These findings, therefore, call for more studies to address basic questions on cell viscosity in physiology and pathology; the proposed approach demonstrated to be a suitable method to study this.

## 4 CONCLUSION

In this paper, the system and method for viscoelasticity measurement using QPI and shear stress is described and tested. Reliability of the proposed viscoelasticity measurement system was tested in several experiments including different cell types and treatments, where the results correspond to the literature and measurements using AFM. The proposed method allows high-throughput measurements during live cell experiments. Processing using parametric deconvolution enables to minimize the effect of the hardware setup on the shear modulus and viscosity estimation. Moreover, we have shown that the proposed approach is suitable for the simultaneous measurement of cell morphology together with cell viscoelastic parameters. Additionally, the proposed method provides a simple approach to measure cell refractive index, which is required for reliable cell height measurement with QPI, where cell height is essential for viscoelasticity calculation.

## AUTHOR CONTRIBUTIONS

T.V., J.G., J.C. and R.K. wrote the manuscript text. T.V., J.G., J.N. and L.C. conducted experiments. T.V. and J.C. wrote the code and conducted analyses on the data. J.N. prepared samples. R.K., V.C., I.P., and M.M. conceived the experimental design and managed the project.

## ACKNOWLEDGMENTS

This work is supported by the Czech Science Foundation, project no. 18-24089S. Computational resources were supplied by the project “e-Infrastruktura CZ” (e-INFRA LM2018140) provided within the program Projects of Large Research, Development and Innovation Infrastructures.

## SUPPLEMENTARY MATERIAL

### 4.1 Cell culture and cultured cell conditions

PC-3 and 22Rv1 cells were used as a model of low- and highly-deformable cells, respectively. Cell lines were purchased from HPA Culture Collections (Salisbury, UK) and were cultured in Ham’s F12 medium with 7 % fetal bovine serum (PC-3) and RPMI-1640 medium with 10 % fetal bovine serum (22Rv1). The RPMI medium was supplemented with antibiotics (penicillin 100 U/ml and streptomycin 0.1 mg/ml). Cells were maintained at 37° C in a humidified (60 %) incubator with 5 % CO_2_ for 2 days prior to exposure to shear stress. The same medium without antibiotics and without fetal bovine serum was used for shear stress induction. The medium was changed twice a week in a routine incubator prior to the measurements. All chemicals were acquired from Merck (Darmstadt, Germany). Cells were further affected by the cytoskeleton-targeting drugs Cytochalasin D (CytD) and docetaxel. Docetaxel treatment was used to stabilise microtubules (23) and hence to increase stiffness and was applied to cells after 48 h post-seeding in a 200 nM concentration for 24 h. CytD is known to interfere with actin polymerisation (21), which leads to a decrease of cell stiffness; it was applied in a concentration of 1 *μ*M for 3 h prior to the measurements.

Before the measurements, the medium for shear stress induction is filtered and preheated to 37° C and its refractive index was determined. Following the connection of fluidics, the system is filled with medium, all visible bubbles are removed, and all components are connected. Patience is given to the connection of the cell cultivation chamber to the fluid system, where no bubbles must occur and the excessive flow (monitored by the flow monitor) caused by the connection must be minimal. With this in mind, a connecting adapter was designed Figure 1c for this purpose.

### 4.2 Atomic force microscopy

To verify the measured shear modulus values, atomic force microscopy was performed. Measurement was carried out on NanoWizard 3 Atomic force microscope (JPK-Bruker, Berlin, Germany), combined with an inverted light microscope (IX-81S1F-3, Olympus, Tokyo, Japan). Force maps were measured in a contact mode using qp-BioT (0.08 N/m) cantilever (Nanosensors) with a 5.73 *μ*m melamine sphere. Mechanical properties were calculated using the DMT model in JPK Data Processing software.

### 4.3 Microfluidic setup optimization

Dependence of the system flow response on the flow circuit components (syringes and tubing) is shown on Figure 8. The curves represent a step response – flow response (measured by the flow meter) to the input step change defined by the syringe pump for various syringes and tubing combinations. These curves clearly demonstrate that the ideal step function is distorted by the nonrigid syringe and tubing and the choice might be critical, particularly when the response time is high (10-30 seconds) or when analyzing the temporal cell response. Surprisingly, the results show that syringes are more important than tubing. Specifically, we found that the usage of glass syringes is crucial for fast system response. Best results were achieved with Fortuna Optima glass syringe 20ml and Darwin Microfluidics PTFE tubing 1/16” / 1/32” (inner diameter/outer diameter).

**Figure 8:**
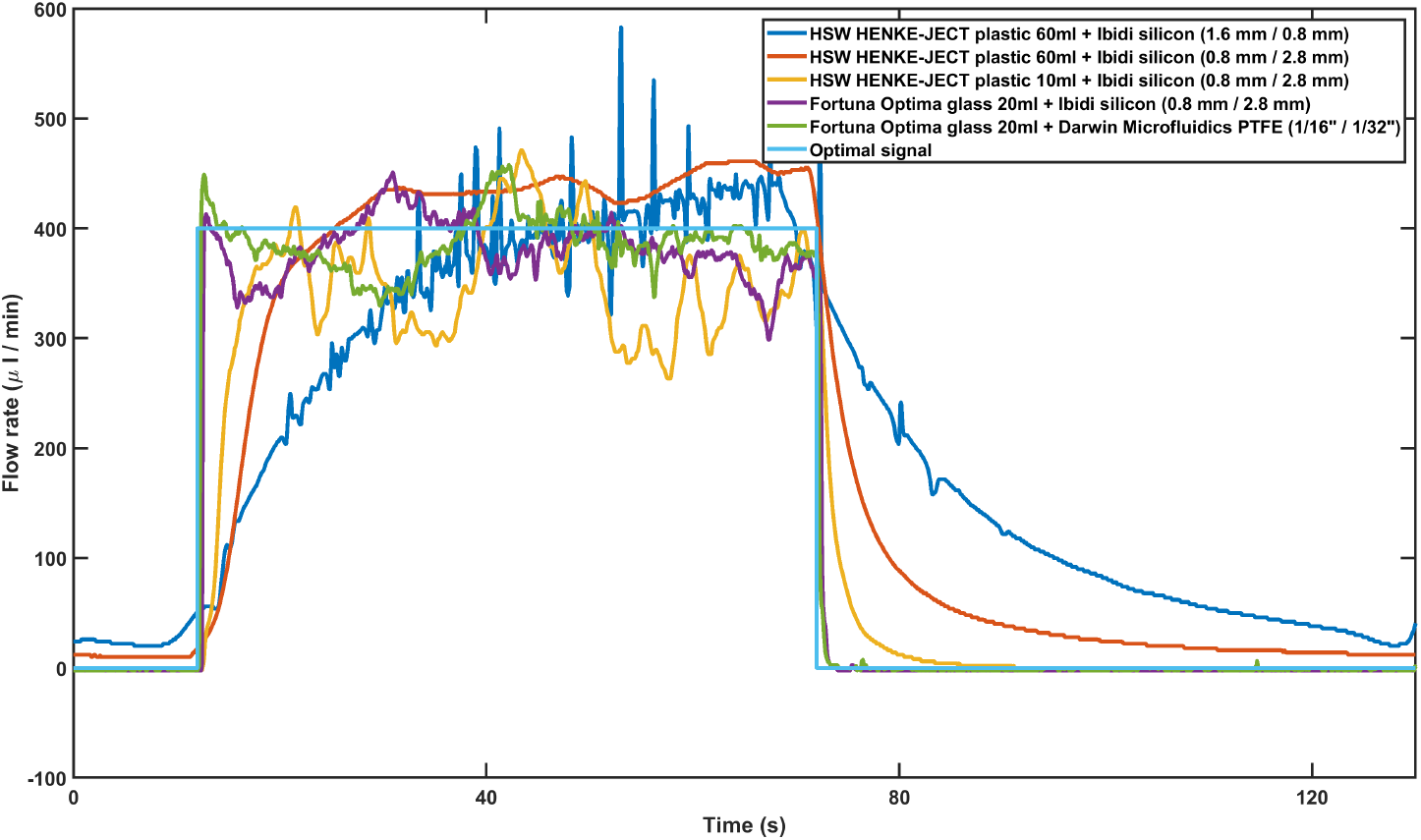
System flow responses to rectangular pulse with various syringes and tubing. Legend shows syringe type + tubing type (inner diameter of tubing/outer diameter of tubing).

### 4.4 Shear stress range optimization

To find the linear range between shear stress and shear modulus (i.e., the range of constant sensitivity), the shear modulus dependence on the shear stress value was measured. In this experiment, a sequence of pulses with increasing shear stress value was applied on PC-3 cells, as shown on Figure 9a. The shear strain value was measured as a difference between local minima and local maxima detected in a 2 second search window located at the beginning and at the end of each pulse in a pulse sequence as measured by flow meter; cell movement was removed from this signal using 3-th order polynomial fitting and subtraction from shear strain signal before this local minima and local maxima measurement. The resulting shear stress-strain dependence is shown in Figure 9b. If we assume a sufficiently long pulse for estimation of the shear strain in steady-state, the equation 3 is reduced to *γ*_∞_ = *τ*_0_/*G*. Value of *γ*_∞_ was measured as the local extreme difference of the shear strain signal. Based on the equation for *γ*_∞_, there should be a linear dependence between stress and strain, where the slope of the resulting line is based on the shear modulus *G* of the cell. However, the expected linear range was limited to 10 Pa and above this value, the saturation of the shear modulus is clearly observable (Figure 9b), thus a larger shear stress is not suitable for shear modulus measurement. As the physiological wall shear stress values in body parts important for cancer metastasis ranges up to 7 Pa in arteries (35) and 9.5 Pa in capillaries (36)); therefore, 0-10 Pa range is suitable for measurement, and 5 Pa was used in the experiments. Accordingly, a high occurrence of cell membrane rupture was observed in cells when exposed to 20 Pa shear stress.

**Figure 9:**
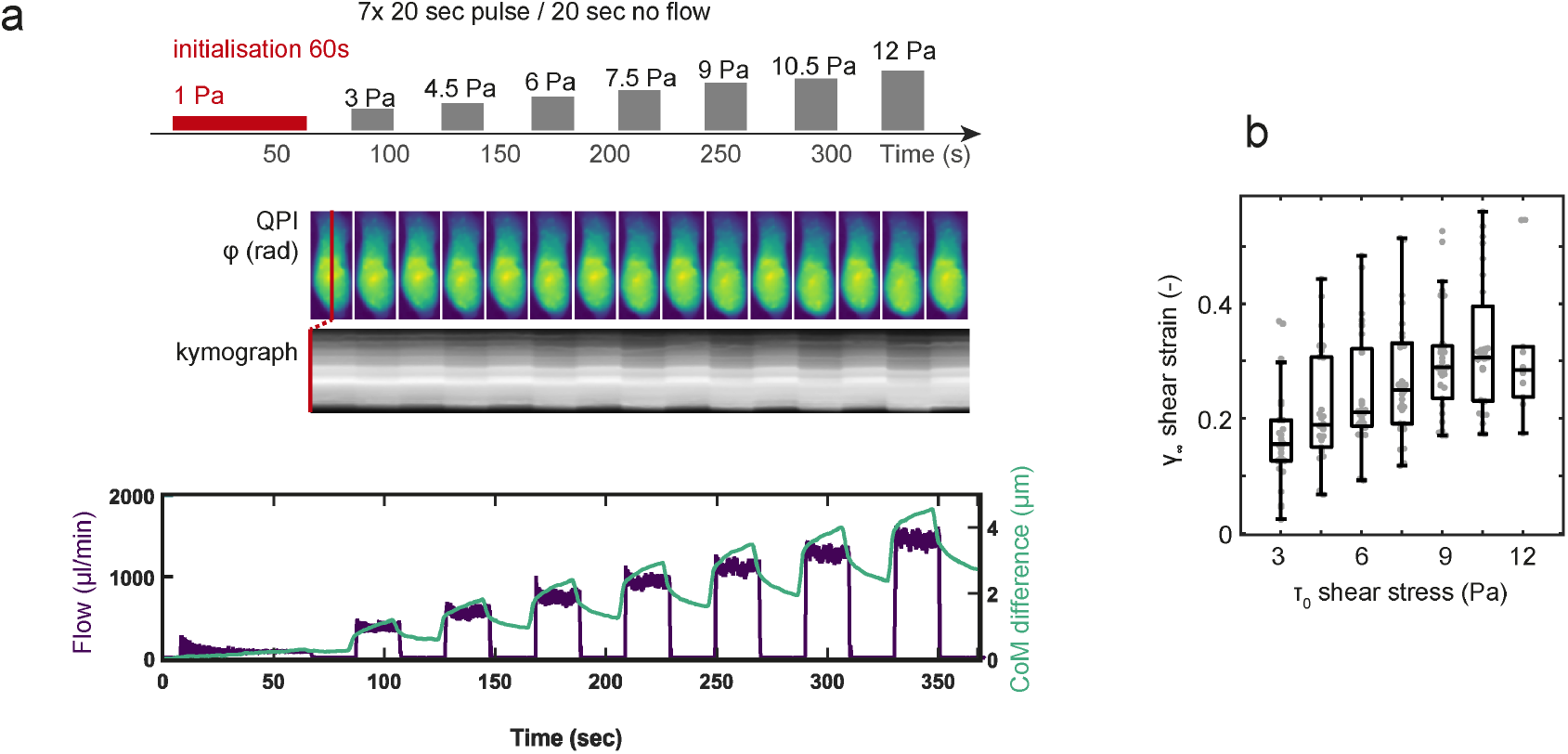
Cell response to applied increasing shear stress sequence. (A) Schematic of experiment with increasing shear stress (top) with kymograph of representative PC-3 cell (middle) and plot of cell centre of mass (CoM) difference over the increasing shear stress sequence together with flow in chamber (bottom). (B) Boxplot representing the dependence of increasing shear stress to measured shear modulus of PC-3 cells with sequence from A)

